# Modelling and Investigating the Interactive Role of Fluid Velocity and Pore Pressure in Load-Induced Osteogenesis

**DOI:** 10.1101/2025.09.22.677695

**Authors:** Himanshu Shekhar, Jitendra Prasad

**Author notes:** Corresponding author **Jitendra Prasad**, Department of Mechanical Engineering, Indian Institute of Technology Ropar, Rupnagar, Punjab, India, 140001. **Himanshu Shekhar** Department of Mechanical Engineering, Indian Institute of Technology Ropar, Rupnagar, Punjab, India, 140001.

## Abstract

Current models propose that osteogenesis occurs in regions of high mechanical stimuli such as strain, fluid velocity, or pore pressure. However, in vivo experiments on mouse tibiae under cantilever loading revealed new bone formation exclusively on the anterolateral side, despite the opposite posteromedial surface experiencing comparable magnitudes of these stimuli. This indicates that individual stimulus magnitude is insufficient and suggests an interactive mechanism among them.

To investigate this, a poroelastic finite element model was developed to quantify the combined effects of load-induced fluid velocity and pore pressure. Tensile loading generated negative pore pressure, stretching osteocyte processes, while compressive loading produced positive pore pressure, compressing them. Since fluid flow exerts drag forces that also stretch osteocytes, the combined effect of flow and negative pressure on the tensile side was hypothesized to enhance mechanotransduction and trigger osteogenesis.

Four potential stimuli were evaluated: dissipation energy density arising from (i) pore pressure, (ii) fluid velocity, (iii) their non-interactive sum, and (iv) their interaction. Comparison with in vivo data showed that only the interactive dissipation energy density accurately predicted both the spatial pattern and rate of new bone formation under high, low, and rest-inserted loading regimes.

These results establish that the interaction between fluid velocity and pore pressure, rather than their independent contributions, governs load-induced osteogenesis. The proposed framework advances the mechanistic understanding of bone adaptation and offers a predictive basis for optimizing mechanical and clinical interventions to promote bone formation and mitigate bone loss.

## 1. Introduction

Nature is renowned for producing structurally optimized structures, and bone is one such example. It continuously adapts and optimizes its microstructure in response to its mechanical loading environment [1]. The optimization is achieved by the coordinated activities of bone-forming cells (osteoblasts) and bone-resorbing cells (osteoclasts) [2]. Another prominent bone cell type, the osteocytes [3] [2] (former osteoblasts), are believed to direct the activities of these two cell types through their intricate lacuno-canalicular network (LCN) embedded in the bone matrix [4]. However, how osteocytes sense mechanical loading/unloading and direct the working cells is still poorly understood.

Over the past few decades, dedicated efforts have been made to decipher the mechanism of bone adaptation. Notably, the pioneering work of Piekarski and Munro[5] contributed substantially to our understanding of load-induced fluid flow within the lacuna-canalicular system (LCS).

Further evidence of load-induced fluid flow was provided by Knoth et al.[6] [7], who used a tracer to confirm its existence within the LCS. After this work, it became well established that load-induced fluid flow is the primary transport mechanism [8], facilitating the movement of biomolecules necessary to maintain bone health.

Estimating fluid velocity induced by mechanical loading remains challenging due to the complexity of the LCS. Very few studies have measured real-time solute transport within the LCS to estimate fluid velocity [9]. To overcome such difficulties, finite element analysis was employed [10] [11] [12]. Subsequently, numerous mathematical models were developed, and fluid flow within the LCS was recognized as a prime stimulus correlating with new bone formation [13] [14] and disuse-induced bone loss [15]. While load-induced fluid flow is well appreciated in the literature, the role of load-induced pore pressure within the LCS in promoting osteogenesis has been largely overlooked.

When bone is mechanically strained, a pressure gradient develops within the LCS that drives fluid flow. However, pore pressure itself cannot be ignored, as it can also radially stretch or compress the osteocyte processes depending on the mechanical loading environment. In vitro studies further confirm the potential role of pore pressure in osteogenesis [16] [17]. Recently, Singh et al. [18] proposed a robust formulation correlating the mineral apposition rate (MAR), a measure of new bone formation, with the dissipation energy density due to load-induced fluid flow and pore pressure. Researchers have modeled cortical bone as a poroelastic material [10] and applied Biot’s theory [19] to estimate pore pressure and fluid velocity, considering the periosteal and endocortical surfaces as impermeable and permeable, respectively.

It is noteworthy that new bone formation has been detected in regions of elevated fluid velocity [18] and pore pressure. However, in a cantilever loading experiment on mouse tibiae, new bone formation occurred at the anterolateral site [20] but not at the posteromedial site, despite both sides exhibiting equal magnitude of peak normal strain (1280 µε), fluid velocity, and pore pressure (Fig. 1). This raises a critical question: Why does this happen?’ Why is the tensile side (anterolateral) more prone to bone formation compared to the compressive side (posteromedial)?

**Figure. 1:**
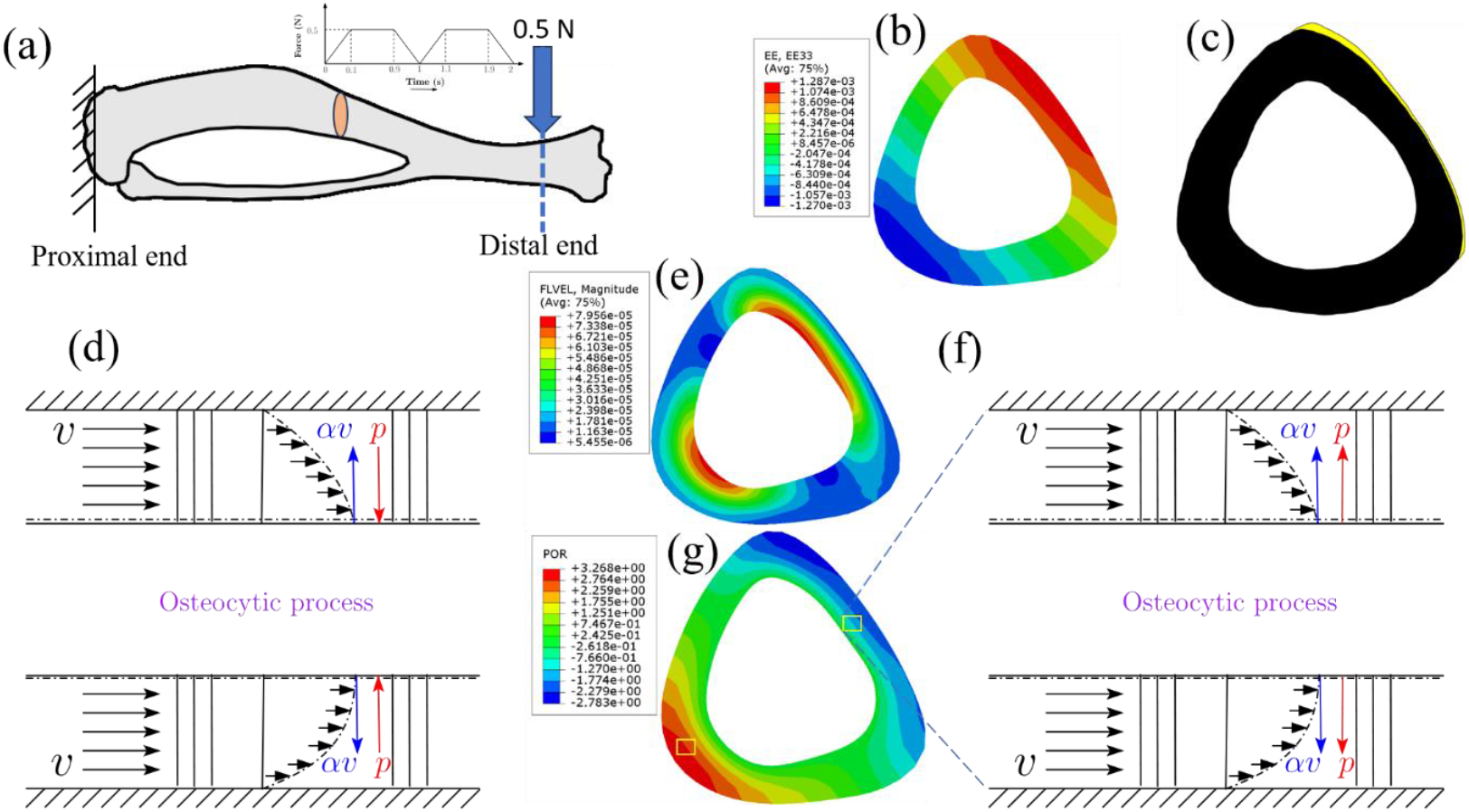
(a) Mouse tibia subjected to cantilever loading of 0.5 N (loading protocol adapted from Srinivasan et al. [22]); (b) corresponding induced normal strain (µε) at the mid diaphysis; (c)induced new bone formation at the anterolateral site; (e) induced fluid velocity (mm/s), where fluid flow tends to expand the cell processes; (g) developed pore pressure (MPa), with positive pressure at the posteromedial site compressing the cell processes and negative pressure at the anterolateral expanding them; (d) magnified view of osteocyte connectivity at the posteromedial site, blue arrow indicates fluid flow action, red arrow indicates pore pressure action; (f) magnified view of osteocyte connectivity at the anterolateral site.

To address this, we first developed a poroelastic model of the mouse tibia, adapting the cantilever loading protocol from the Srinivasan et al. [20]. Interestingly, while fluid velocity was similar at both sites, the sign of pore pressure was opposite. Tensile strain at the anterolateral surface induced negative pore pressure (Fig. 1(g)), promoting radial stretching of the osteocyte cell wall (Fig. 1(f)), similar to the effect of fluid flow [21]. In contrast, compressive strain at the posteromedial site induced positive pore pressure, which acted opposite to the fluid flow on the cell processes (Fig. 1(d)). Thus, the net algebraic effect at the tensile site was significant and promoted osteogenesis. This observation motivates the present work. We hypothesized that the combined effect of fluid flow and pore pressure is greater at the tensile site, thereby driving new bone formation.

To test this hypothesis, we examined four cases: (1) the role of dissipation energy density induced by pore pressure alone in new bone formation; (2) the dissipation energy density based on fluid velocity as a stimulus for osteogenesis; (3) the combined effect of dissipation energy densities due to fluid velocity and pore pressure; and (4) the dissipation energy density arising from the interaction between fluid velocity and pore pressure. We present a robust methodology to capture the role of dissipation energy at osteocytes resulting from the combined effects of load-induced fluid flow and pore pressure. The developed mathematical model was validated using existing cantilever loading-induced new bone formation data from the literature [20]. The model’s accuracy was evaluated using statistical tests, viz., Student’s t-test and Watson U^2^ test. The t-test compared the in-vivo bone formation rate per unit bone surface (BFR/BS) with the model-predicted values, while the Watson U^2^ test compared the in-vivo site-specific distribution of new bone formation with that predicted by the model.

The paper is structured as follows: Section 2 outlines the methodology, Section 3 presents the results, Section 4 discusses the findings, and Section 5 provides the conclusions.

## 2. Methodology

### 2.1. Experimental Data

The in vivo data for the cantilever loading protocol, including the corresponding BFR/BS values and site-specific new bone formation distributions, were adapted from the study by Srinivasan et al. [20]. Three loading regimes were considered, as detailed below.

#### 2.1.1 Low Magnitude

The low-magnitude loading protocol was conducted on 10-week-old C57BL/6J female mice (*n* = 6). A trapezoidal waveform with a strain rate of 0.01/s and a peak load of 0.25 N was applied for 5 consecutive days (100 cycles per day). This loading induced a peak normal strain of 650 µε at the mid-diaphysis of the mouse tibia. The corresponding BFR/BS value was 0.011 ± 0.013 *μm*^3^/*μm*^2^/*day*.

#### 2.1.2 Low Magnitude with Rest-Insertion

The low-magnitude loading protocol with 10 s rest insertion was also conducted on 10-week-old C57BL/6J female mice (*n* = 6). A trapezoidal waveform with a strain rate of 0.01/s and a peak load of 0.25 N was applied for 5 consecutive days (10 cycles per day), with a 10 s rest period between successive loading cycles. This induced a peak normal strain of 650 µε at the mid-diaphysis. The corresponding BFR/BS value was 0.095 ± 0.071*μm*^3^/*μm*^2^/*day*.

#### 2.1.3 High Magnitude

The high-magnitude loading protocol was performed on 10-week-old C57BL/6J female mice (*n* = 6). A trapezoidal waveform with a strain rate of 0.02/s and a peak load of 0.5 N was applied for 5 consecutive days (100 cycles per day). This loading induced a peak normal strain of 1300 µε at the mid-diaphysis. The corresponding BFR/BS value was 0.2238 ± 0.1221*μm*^3^/*μm*^2^/*day*.

### 2.2 Loading and Boundary Conditions

The analysis was focused on the mid-diaphysis of the mouse tibia; performing a poroelastic analysis of the entire tibia would be computationally expensive. The bone was idealized as a prismatic bar, 3 mm in length, with a cross-section similar to that of the mid-diaphysis of a C57BL6J mouse tibia studied by Srinivasan et al. [20]. In the literature, the periosteal surface is generally considered impermeable, while the endocortical surface is permeable[18], respectively. Accordingly, these conditions were implemented as zero-flow and zero-pressure boundary conditions, respectively.

Three different trapezoidal loading waveforms, low magnitude (Fig.2(d)), low magnitude with rest insertion (Fig. 2 (e)), and high magnitude (Fig. 2(f)), were adapted from Srinivasan et al. [22] and applied in the finite element analysis to compute the strain, fluid velocity, and pore pressure (see Section 2.3).

**Figure. 2:**
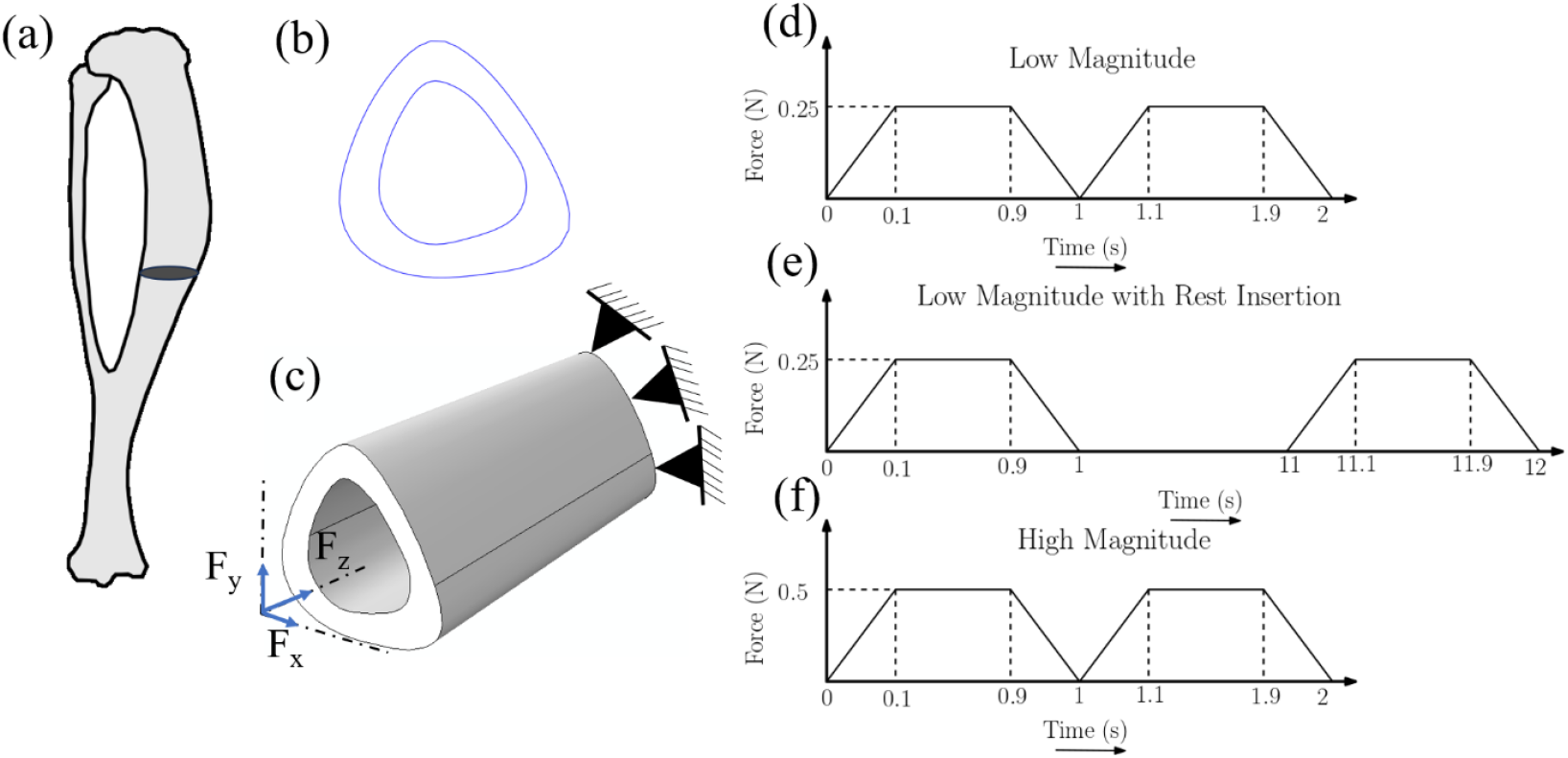
(a) Tibia and fibula of a mouse, with the cross-section of interest highlighted; (b) cross-section of interest as studied by Srinivasan et al. [22]; (c) prismatic bar of 3mm length created for poroelastic analysis, with a cross-section similar to that used in the experiment. One end of the prismatic bar was fixed, while the other end was loaded in all three directions (x, y, and z) to induce strains similar to those reported by Srinivasan et al.[22]. The loading waveforms were also adapted from the same study: (d) low magnitude; (e) low magnitude with 10 s rest between cycles; (f) high magnitude.

### 2.3 Finite Element Analysis

Cortical bone consists of a solid matrix and pores through which biofluid passes to maintain homeostasis. These porosities exist at different length scales, such as vascular, lacuna-canalicular, and collagen-fiber levels. Dynamic load-induced fluid flow and pore pressure developed within the lacuna-canalicular porosities are primarily considered stimuli that promote osteogenesis. Therefore, only LCN porosity was considered in the present poroelastic analysis, and the corresponding permeability value was adapted from earlier studies on mouse tibia (Table 1) [23] [24]. The behavior of such a poroelastic material under loading was analyzed using Biot’s theory of poroelasticity, and the finite element analysis was conducted in ABAQUS [25]. The poroelastic material properties were taken from earlier mouse tibia studies [15] [26] and are listed in Table 1.

**Table 1:** Poroelastic material properties used in the simulations (adapted from Kumar et al.[26][15] except permeability value from Zhou et al.[24] [27])

| Properties | Magnitude | Unit |
| --- | --- | --- |
| Young's modulus of bone | 12 | GPa |
| Fluid bulk modulus | 2.3 | GPa |
| Solid bulk modulus | 17 | GPa |
| Porosity | 0.05 |  |
| Poisson's ratio | 0.3 |  |
| Intrinsic permeability (k) | $3.79 \times 10^{-21}$ | $m^2$ |

The prismatic bar, 3 mm in length, was meshed with 15,120 linear hexahedral elements after convergence analysis. ABAQUS provides the element type C3D8RP (continuous 3-dimensional 8-noded reduced integration pore pressure) for analyzing poroelastic behavior under loading. These elements have four degrees of freedom per node: three displacement components (*u_x_, u_y_*, and *u_z_*) in the x, y, and z directions, respectively, and pore pressure as the fourth. Darcy law, which relates fluid velocity to the pressure gradient, was used to compute fluid velocity at the integration points. Further details of the constitutive relation can be found in the earlier study [15].

One end of the meshed prismatic bar was fixed, and the other end was loaded in all three directions to induce strains similar to those in the experimental study. The obtained strain pattern and corresponding peak strain values were verified and found to agree closely with the experimental values. Load-induced strain (EE33), fluid velocity, and pore pressure are shown in Fig. 3. Notice that strain magnitudes at the tensile and compressive sides of the cross-section are approximately the same: 634 *με* and −628 *με*, respectively, for the low-magnitude loading protocol (including the case with rest-insertion); and 1287 *με* and −1270 *με*, respectively, for the high-magnitude loading protocol. Despite these similar strain magnitudes, new bone formation was experimentally observed only on the tensile side (Fig. 1(c)).

**Figure. 3:**
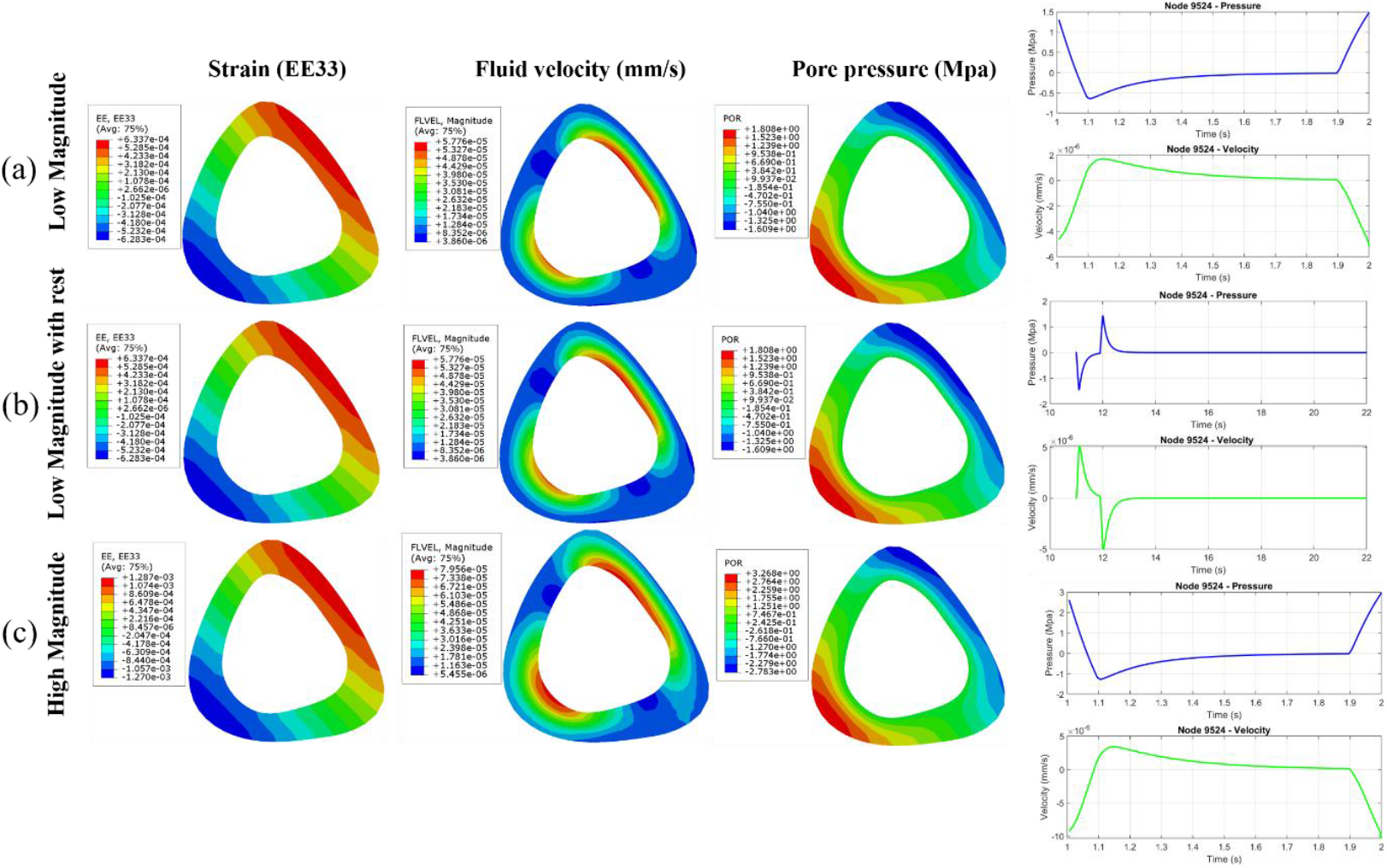
Contour plots from finite element analysis under three loading protocols: (a) low-magnitude loading showing strain (µε), fluid velocity (mm/s), pore pressure (MPa), and pressure-velocity vs. time at node 9524 (tensile side of the cross-section); (b) low-magnitude loading with 10 s rest between cycles showing strain (µε), fluid velocity (mm/s), pore pressure (MPa), and pressure-velocity plot vs. time at node 9524; (c) high-magnitude loading showing strain (µε), fluid velocity (mm/s), pore pressure (MPa), and pressure-velocity vs. time at node 9524.

### 2.4 Derivation of Load-induced Dissipation Energy Density

We computed the load-induced fluid velocity and pore pressure with respect to the applied loading protocol at the osteocyte level. These load-induced stimuli act directly on osteocytes and their cell processes. Fluid flow induces radial stretching, tending to expand the cell processes [28]. In contrast, the effect of pore pressure is sign-dependent: negative pressure (observed on the tensile strain side at the anterolateral surface of the mid-diaphysis) stretches the cell processes (Fig. 4), similar to fluid flow; whereas positive pore pressure compresses the cell processes, opposing the action of fluid flow.

**Figure. 4:**
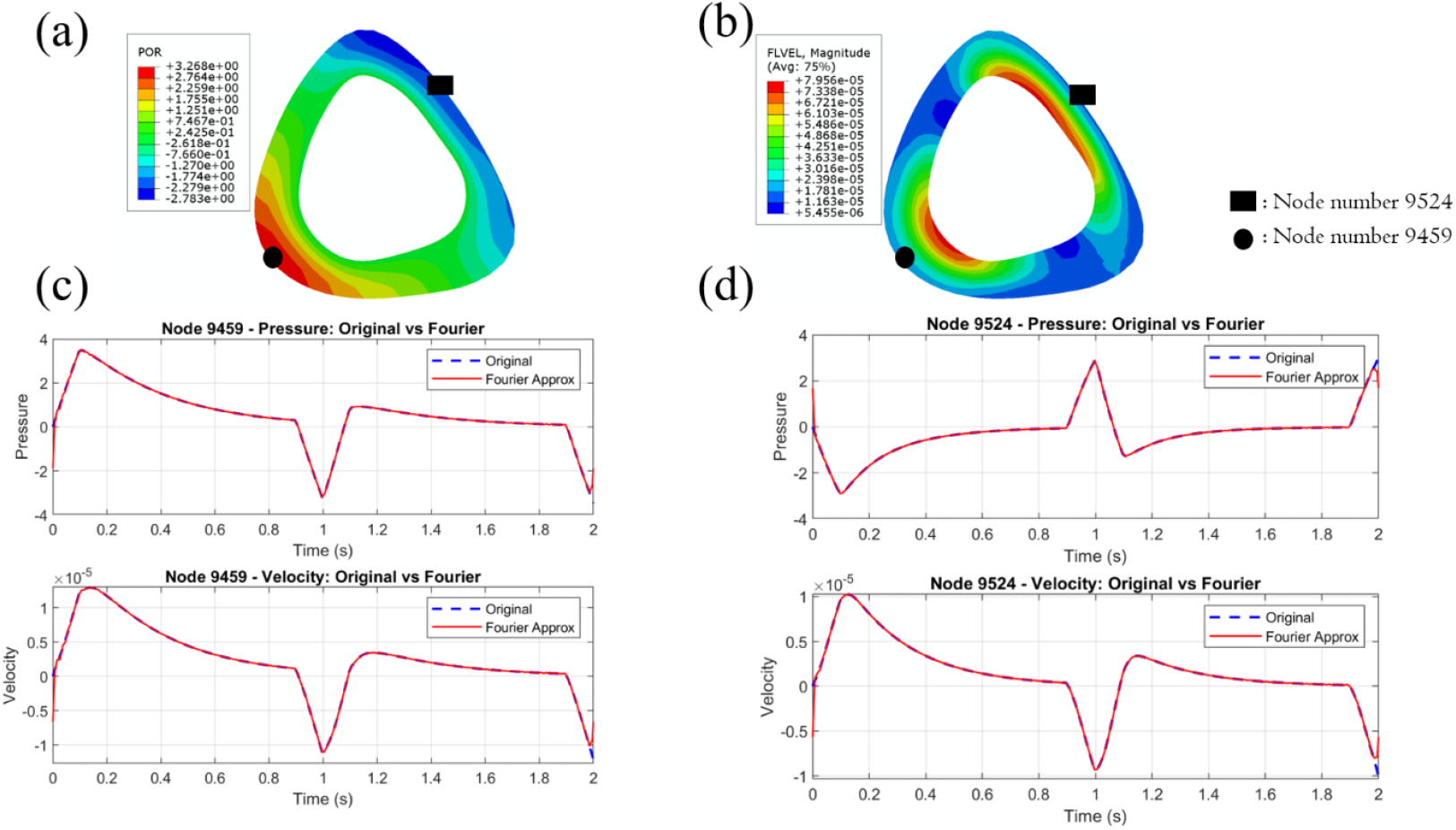
Interaction of fluid velocity and pore pressure:(a) contour of cantilever loading-induced pore pressure (black filled rectangle denotes node 9524 at the anterolateral surface, whereas black filled circle represents node 9459 at the posteromedial surface) at the mid-section; (b) contour of fluid velocity (mm/s); (c) plot of pore pressure and fluid velocity with respect to time. It is evident that the effect of fluid flow attempts to expand the cell processes, while a positive pore pressure attempts to compress the cell processes at the posteromedial site; (d)plot of pore pressure and fluid velocity with respect to time. Here, the effects of both are additive in nature, and hence, the net effect attempts to expand the cell processes.

The effect of fluid flow can be described using viscous damping, where the damping force (F) is directly proportional to the velocity [21] [29].

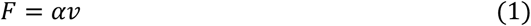

where *α* is a proportionality constant. The net radial stretch at a point can therefore be expressed as: *σ*(*t*) = −*p*(*t*) + *αv*(*t*).

Osteocytes and their processes can be modeled as viscoelastic materials under the influence of these stimuli, which leads to viscoelastic dissipation. The corresponding dissipation energy density has been adopted as a stimulus for new bone formation.

We idealized viscoelastic dissipation through the Kelvin-Voigt model (Fig. 5(b)), which represents the material response as a spring and a damper connected in parallel:

**Figure. 5:**
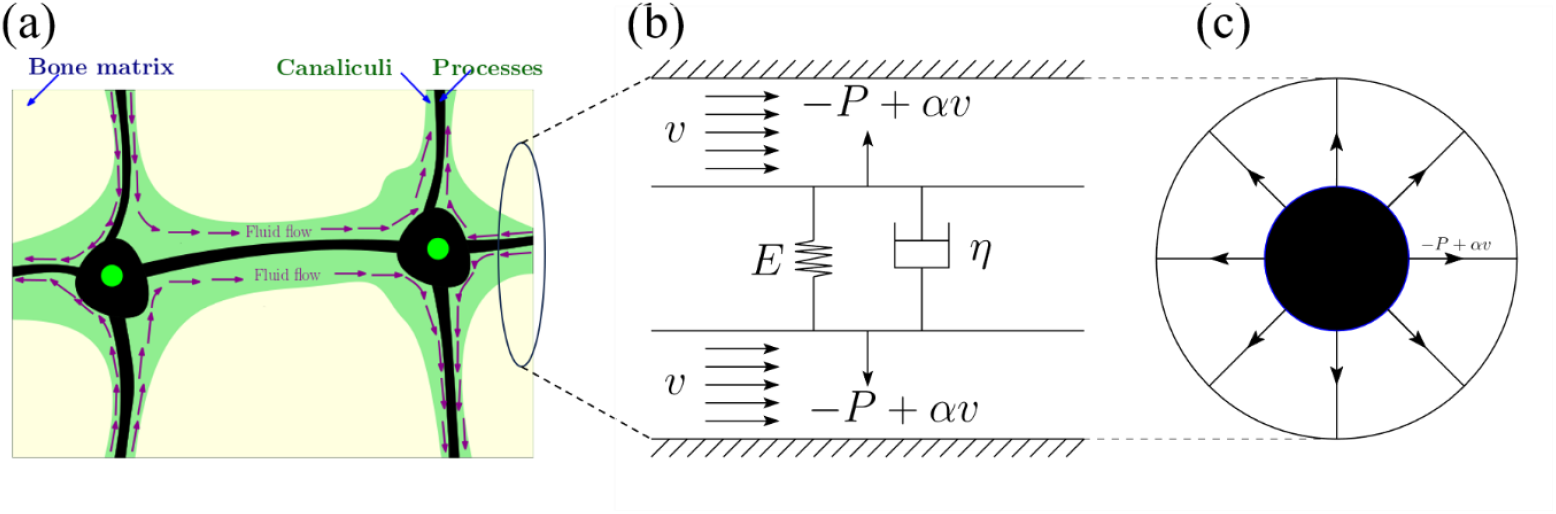
(a) Idealized lacuna-canalicular system, with the arrow indicating load-induced fluid flow in the pericellular space; (b) osteocytes and their processes modeled as a viscoelastic material under the influence of the net stretching effect, where E, and դ represent the elastic and viscous terms, respectively; (c) left-hand side view of the cross section of a canaliculus, where the black filled circle represents the cell process and the radially outward lines depict the tethering fibers.

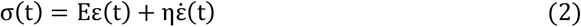

where E is the elastic modulus, η is the viscosity, ε(t) is strain, and 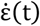 is strain rate. Here, *σ*(*t*) is a time-dependent function, which can be expressed as a Fourier series:

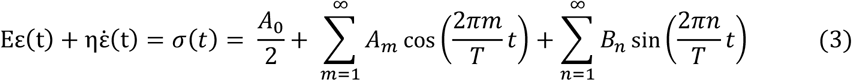

where, *A*_0_, *A*_*m*_, and *B*_*n*_ are constants, and *m* and *n* are harmonic numbers. The viscoelastic dissipation energy density per unit cycle of loading is:

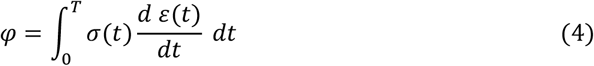

Since *σ*(*t*) is known (as a function of pore pressure and fluid velocity), *ε*(*t*) can be easily calculated from Eq. (2), with boundary condition *ε*(*t* = 0) = 0:

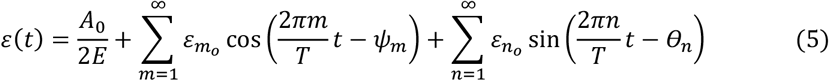

and

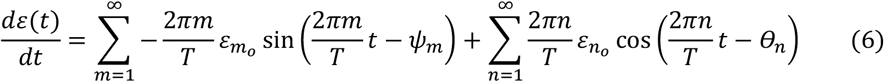

Using Eq. (4), dissipation energy density per cycle can be written as:

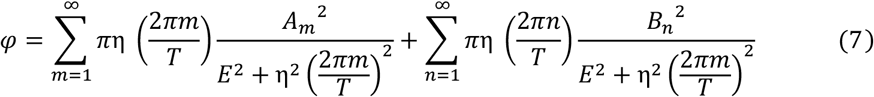

where, *A*_*m*_ and *B*_*n*_ are:

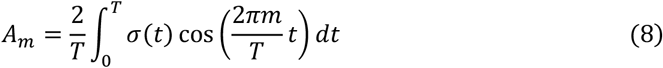

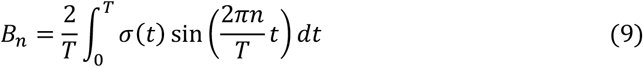

Since *σ*(*t*) = −*p*(*t*) + *αv*(*t*), the Fourier expansions yield (with *T*=*T*_0_):

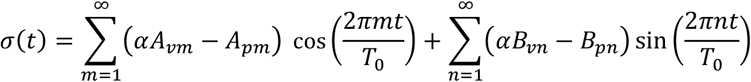

Hence, dissipation energy density becomes:

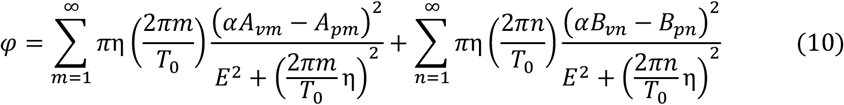

This can be decomposed as:

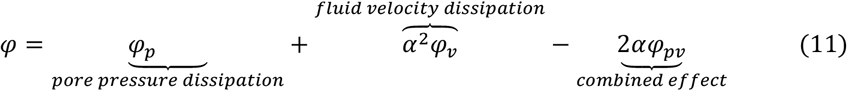

where,

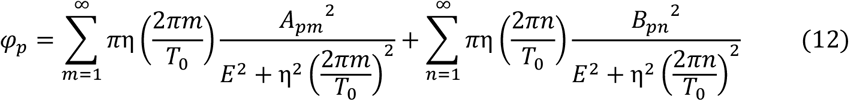

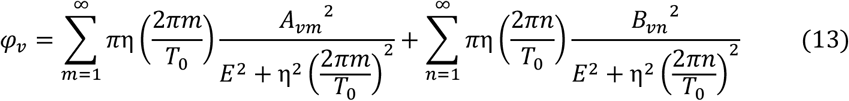

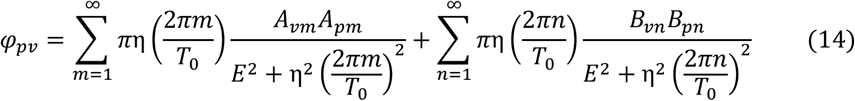

with Fourier coefficients:

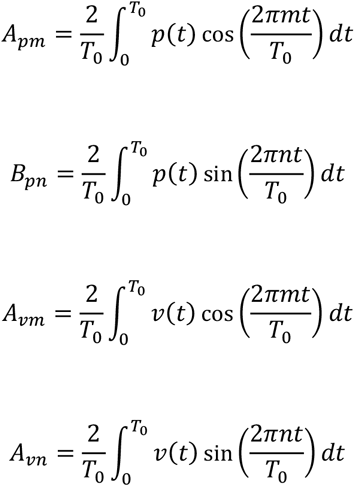

### 2.5 Zone of Influence

Each osteocyte contributes to osteogenesis; however, its influence decreases as the distance from the osteocyte increases, and is greater when the target location is nearby. This non-local behavior of osteocytes has been modeled in the literature using different approaches. Prasad and Goyal [30] employed a diffusion-based model, while Singh et al. [18], Kumar et al.[26], and Shekhar et al. [15] used the zone of influence (ZOI) concept. In the present study, the ZOI approach was adopted to average the stimulus (dissipation energy density) from each osteocyte, as illustrated in Fig. 6(a). A weightage average scheme was applied such that nodes farther from the target node contributed less, while closer nodes contributed more. The exponential weighting function *w* = *e*^−5(|*x*|*/r*)^ was adopted from earlier studies [26], where r is the radius of the ZOI and |x| is the distance between the target node and a node within the ZOI. The value of r was taken as 170 µm, corresponding to the mean cortical thickness at the mid-diaphysis of the mouse tibia (average of maximum and minimum cortical thickness).

**Figure. 6:**
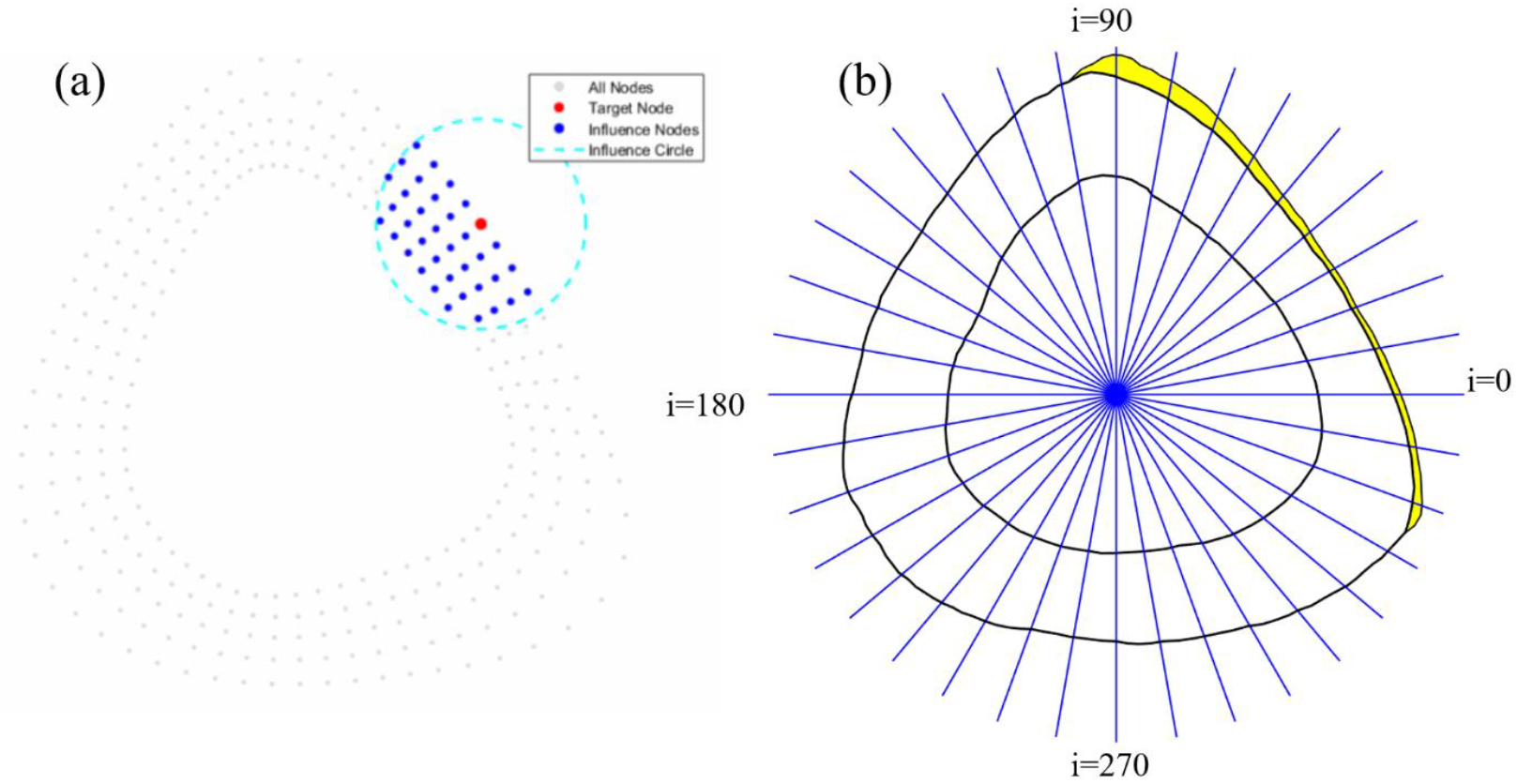
(a) Averaging of the stimulus using the zone of influence (ZOI). Gray nodes represent the overall bone cross-section, the red node indicates the target node where the stimulus is computed, and blue nodes denote the contributing nodes influencing the target node. (b) Computation of experimental MAR at each i^th^ node of interest using an in-house code (Yellow color indicates new bone formation).

### 2.6 Computation of Mineral Apposition Rate

The site-specific new bone formation is commonly quantified using the parameter mineral apposition rate (MAR). Previous studies correlated MAR with different stimuli such as load-induced dissipation energy density, strain rate, and strain energy density. However, no robust or universally accepted relationship has been established. Recently, Singh et al. [18] proposed a formulation suggesting that MAR is directly proportional to the square root of the dissipation energy density minus a threshold value, with osteogenesis is triggered only when the stimulus exceeds this threshold. Notably, this concept holds not only for new bone formation, but also for disuse-induced bone loss [23].

This relation can be written as:

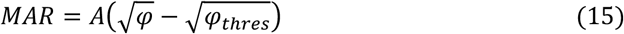

where *A* is the remodeling rate. Hence, using Eqs. (10) and (11), *MAR* can be expressed as:

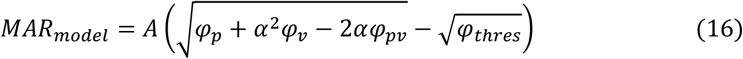

Here, *A*, α, and *φ*_*thres*_ are constants to be determined. *MAR* (mm^3^/mm^2^/da*y*) is defined as the total new bone volume formed per unit bone surface per unit time i.e., the new bone thickness added at each point on the cross-section.

For computational convenience, the periosteal surface was divided into 360 segments, and the corresponding MAR value was computed at each i^th^ point (the common point between the two segments), as shown in Fig. 6(b):

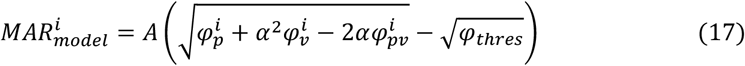

where, 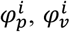, and 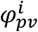 are known from Eqs. (10) and (11). The constants *A*, α, and *φ*_*thres*_ were determined by comparing with experimental new bone formation data reported in the literature [20]. The experimental 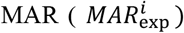 was computed at 360 points (Fig. 6(b)) using an in-house MATLAB code. The constants were obtained by minimizing the squared error:

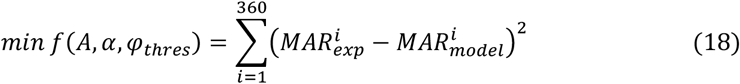

The minimization problem was solved using the Levenberg-Marquardt algorithm in MATLAB, which fits the model Eq. (17) to the experimental 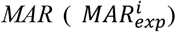 of the high-magnitude loading protocol and computes the optimal value of *A*, α, and *φ*_*thres*_. The obtained optimal values were:

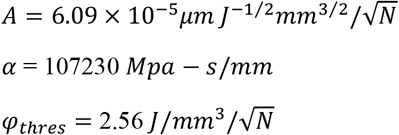

These constants were subsequently used to predict MAR for the other two loading protocols, i.e., low magnitude and low magnitude with rest insertion. Model predictions were evaluated using two statistical tests: Student’s t-test and Watson’s U^2^ test. The Student’s t-test compared the model bone formation rate (BFR/BS) with the experimental BFR value, while the Watson’s U^2^ test compared the site-specific new bone distribution.

### 2.7 Statistical Analysis

The new bone formation area per unit time was calculated by integrating *MAR* over the periosteal circumference. The bone formation rate per unit bone surface (BFR/BS) was then obtained by dividing the new bone formation area per unit time by the total perimeter of the periosteal surface of the mouse tibia.

A one-sample, two-tailed Student’s t-test was used to compare the experimental BFR/BS values reported in the literature with those predicted by the model. In addition, a circular goodness-of-fit test, viz., Watson’s U^2^ test [31] was employed to compare the experimental site-specific distribution of new bone formation with that predicted by the model. A model is considered to fit the experimental data only if *p* > 0.05 for both the Student’s t-test and Watson’s U^2^ test.

The null hypothesis states that none of the four dissipation energy density cases — (i) due to pore pressure only (*φ*_*p*_), (ii) fluid velocity only (*α*^2^*φ*_*v*_), (iii) their non-interactive combination (*φ*_*p*_ + *α*^2^*φ*_*v*_), and (iv) their combination (*φ* = *φ*_*p*_ + *α*^2^*φ*_*v*_ − 2*αφ*_*pv*_) — can reproduce the experimental results where cantilever loading of the murine tibia leads to new bone formation on the tensile side of the cross-section but not on the compressive side. The alternative hypothesis is that only the interactive combination of pore pressure and fluid velocity can correctly model this asymmetric bone formation.

## 3. Results

### 3.1 Dissipation Energy Density Due to Pore Pressure Alone as a Stimulus

**High Magnitude:** Initially, the dissipation energy density induced by pore pressure (*φ*_*p*_) was used as the sole stimulus to promote new bone formation. The site-specific new bone formation predicted by the model was compared with the in vivo distribution (Fig. 7(c)). The Watson U^2^ test indicated that the model-predicted distribution was significantly different from the in-vivo distribution (p= 0.025, Watson U^2^ test). Hence, pore pressure alone is not sufficient to predict new bone formation. The BFR/BS values were also compared (Fig. 7(d)); however, no significant difference was observed (p = 0.92, Student’s t-test).

**Figure. 7:**
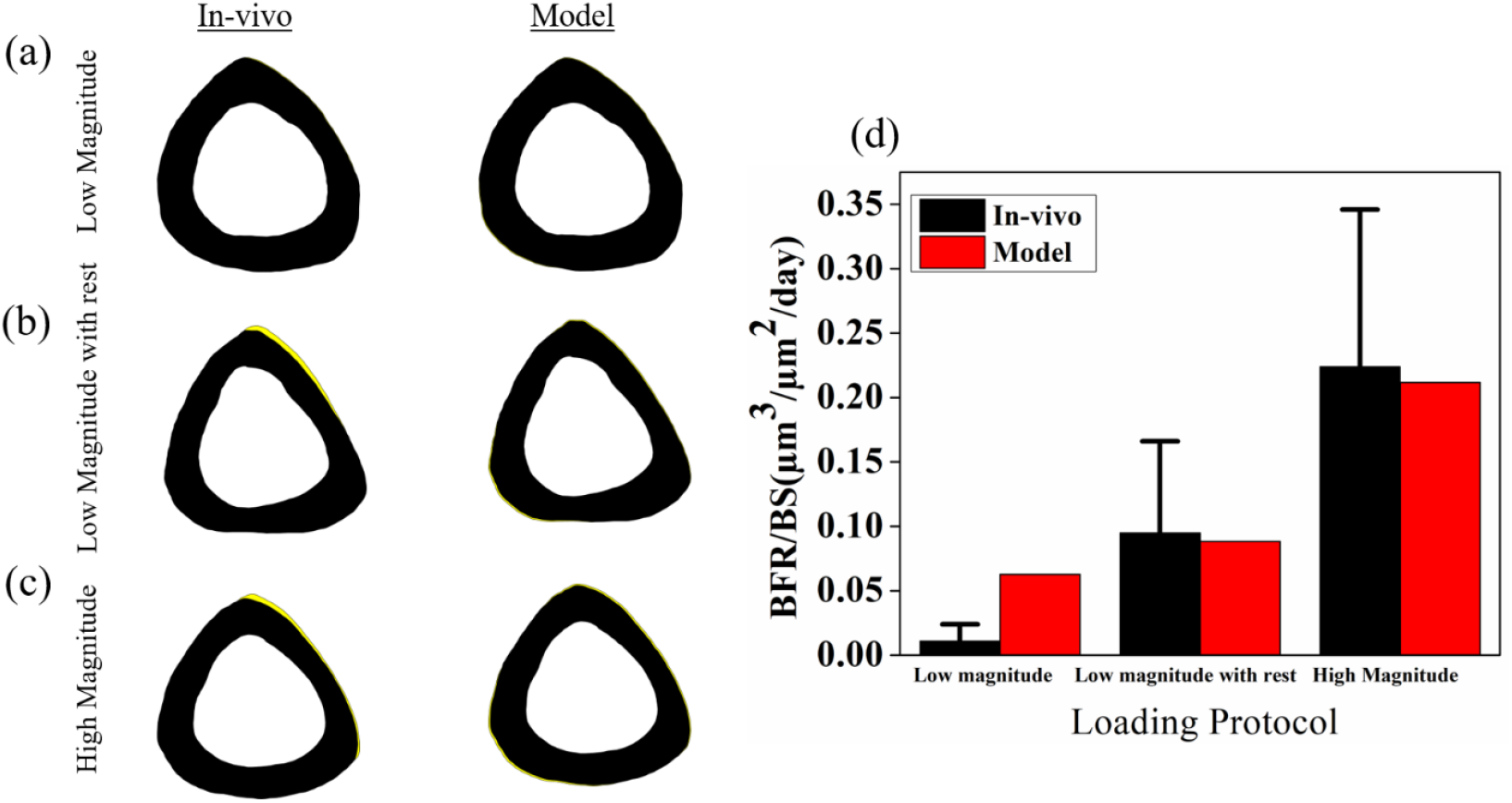
Comparison of experimental new bone formation with model predictions when dissipation energy density induced by pore pressure alone (*φ*_*p*_) utilized as a stimulus: (a) site-specific new bone formation under the low-magnitude loading protocol (yellow indicates new bone formation); (b) site-specific new bone formation under the low-magnitude loading protocol with rest insertion; (c) site-specific new bone formation under the high-magnitude loading protocol; and (d) comparison of experimental and model-predicted BFR/BS values.

**Low Magnitude:** A low-magnitude loading protocol was also analyzed, considering pore pressure-induced dissipation energy density alone (*φ*_*p*_) as the mechano-transductive stimulus. Both the site-specific bone distributions (p = 0.003, Watson U^2^ test; Fig. 7(a)) and BFR/BS values (p = 0.04, Student’s t-test; (Fig. 7(d)) were found to be significantly different from those observed in vivo.

**Low Magnitude with Rest Insertion:** When pore pressure alone was considered as the stimulus (*φ*_*p*_) under the rest-insertion waveform with low magnitude (Fig. 6(b)), the model failed to predict the site-specific bone distribution (p = 3 × 10^−5^, Watson U^2^ test). However, no significant difference was observed in the BFR/BS values (p = 0.62, Student’s t-test).

### 3.2 Dissipation Energy Density Due to Fluid Velocity Alone as a Stimulus

**High Magnitude:** In the second test case, the dissipation energy density due to fluid flow was utilized solely (*α*^2^*φ*_*v*_) as a stimulus for osteogenesis. The site-specific distributions and BFR/BS values were compared (Fig. 8). The site-specific distribution predicted by the model was found to be significantly different from the in vivo distribution (p = 0.036, Watson U^2^ test; Fig. 8(c)). Furthermore, the BFR/BS values were compared through a bar graph (Fig. 8(d)); however, no significant difference was observed (p = 0.72, Student’s t-test).

**Figure. 8:**
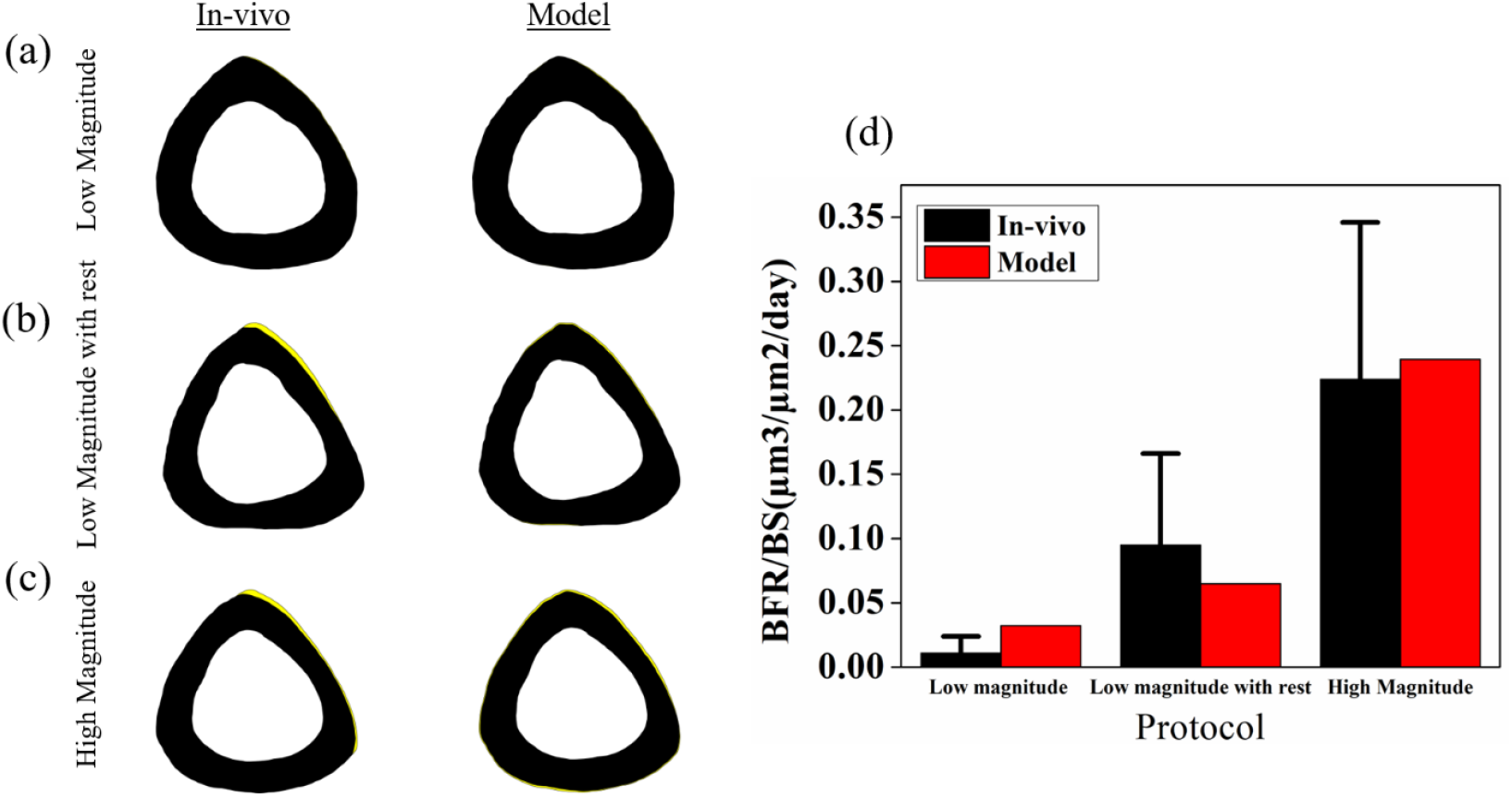
Comparison of experimental new bone formation with model prediction when dissipation energy density induced by interstitial fluid velocity alone (*α*^2^*φ*_*v*_) was utilized as the stimulus: (a) site-specific new bone formation under the low-magnitude loading protocol (yellow indicates new bone formation); (b) site-specific new bone formation under the low-magnitude loading protocol with rest insertion; (c) site-specific new bone formation under the high-magnitude loading protocol; (d) comparison of experimental and model-predicted BFR/BS values.

**Low Magnitude:** The site-specific new bone distributions under the low magnitude loading protocol were also compared (Fig. 8(a)). The model-predicted distribution was significantly different from the in vivo distribution (p = 0.046, Watson U^2^ test). However, BFR/BS values were not significantly different (p = 0.30, Student’s t-test).

**Low Magnitude with Rest Insertion:** The dissipation energy density induced by fluid flow alone (*α*^2^*φ*_*v*_) was not sufficient to replicate the in vivo site-specific bone distribution under the low magnitude loading protocol with rest insertion (p = 0.011, Watson U^2^ test (fig. 8(b)). However, the model-predicted BFR/BS value was not significantly different from the in vivo BFR/BS value (p = 0.69, Student’s t-test).

### 3.3 Dissipation Energy Density Due to the Combination of Fluid Velocity and Pore Pressure Without Interaction as a Stimulus

**High Magnitude:** The third test case incorporated dissipation energy density arising from both fluid velocity and pore pressure without interaction (*φ*_*p*_ + *α*^2^*φ*_*v*_). This combination was first used to predict the high-magnitude loading protocol. The site-specific comparison is shown in Fig. 9(c), where the model-predicted distribution was found to be significantly different from the in vivo distribution (p = 0.035, Watson U^2^ test). The BFR/BS values were compared using a bar chart (Fig. 9(d)), and no significant difference was observed (p = 0.71, Student’s t-test).

**Fig. 9:**
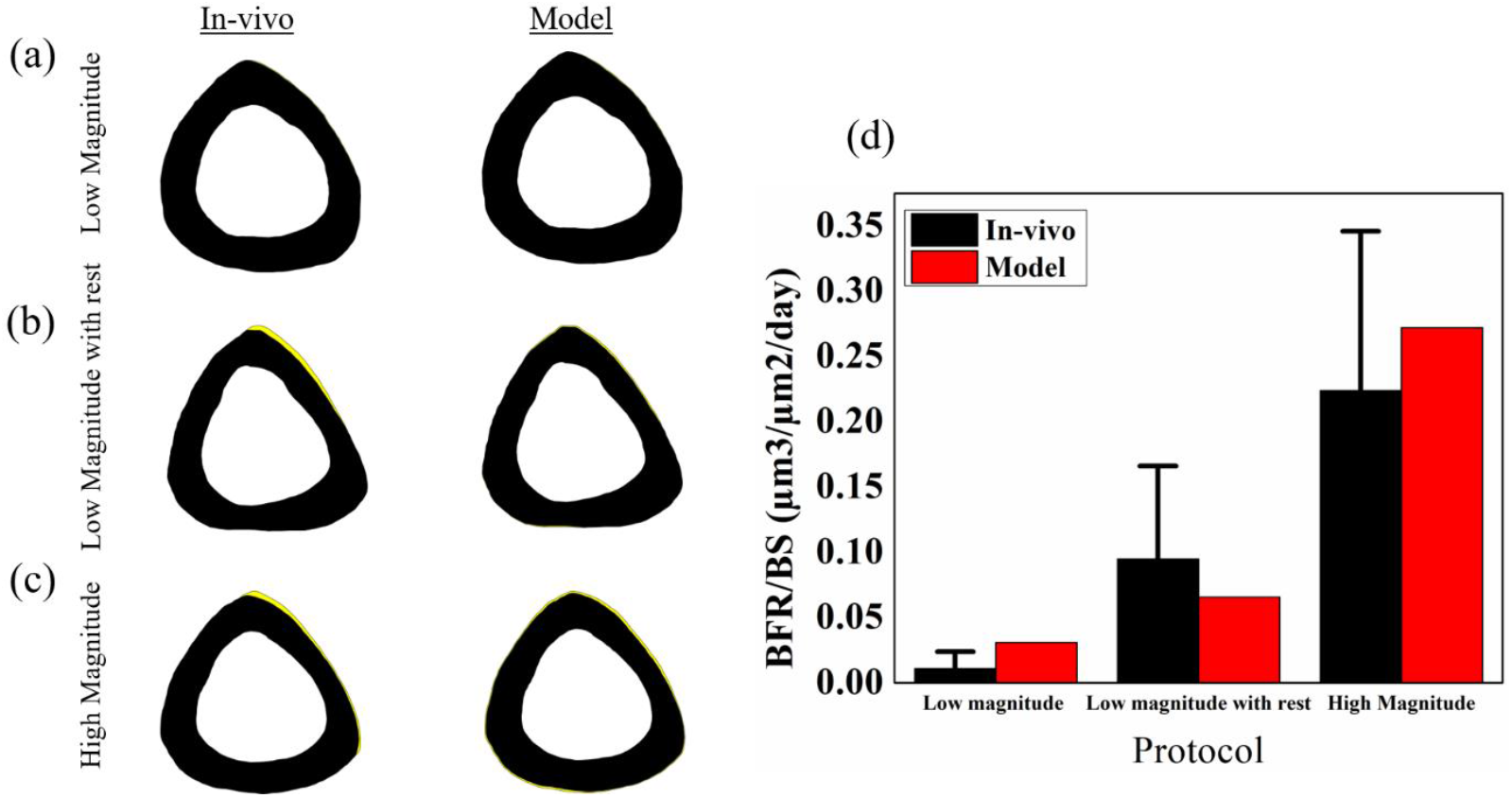
Comparison of experimental new bone formation with model prediction when dissipation energy density induced by both pore pressure and fluid velocity without interaction (*φ*_*p*_ + *α*^2^*φ*_*v*_) was utilized as the stimulus: (a) site-specific new bone formation under the low-magnitude loading protocol (yellow indicates new bone formation); (b) site-specific new bone formation under the low-magnitude loading protocol with rest insertion; (c) site-specific new bone formation under the high-magnitude loading protocol; (d) comparison of experimental and model-predicted BFR/BS values.

**Low Magnitude:** The low-magnitude loading protocol was also predicted through this combination (Fig. 9(a)); however, the site-specific distribution was significantly different from the in vivo distribution (p = 0.04, Watson U^2^ test). The BFR/BS values were also compared (Fig. 9(d)), and no significant difference was found (p = 0.33, Student’s t-test).

**Low Magnitude with Rest Insertion:** For the low-magnitude protocol with rest insertion, the site-specific new bone distribution was again found to be significantly different from the in vivo distribution for this case also (p = 0.007, Watson U^2^ test; Fig. 9(b)). However, the BFR/BS values were not significantly different (p = 0.70, Student’s t-test).

### 3.4 Dissipation Energy Density Due to the Interaction of Fluid Velocity and Pore Pressure as a Stimulus

**High Magnitude:** In this case, the dissipation energy density resulting from the interaction between the fluid velocity and pore pressure (*φ* = *φ*_*p*_ + *α*^2^*φ*_*v*_ − 2*αφ*_*pv*_) was utilized as a mechano-transductive stimulus. The BFR/BS values were compared using a bar chart (Fig. 10(d)), and the model-predicted BFR/BS (0.2354 *μm*^3^/*μm*^2^/*day*) was not significantly different from the experimental BFR/BS value (0.2238 ± 0.1221 *μm*^3^/*μm*^2^/*day*, p =0.87, Student’s t-test). The experimental site-specific new bone formation (Fig. 10(c)) was also compared with the model-predicted distribution, and no significant difference was observed (p = 0.70, Watson U^2^ test).

**Figure. 10:**
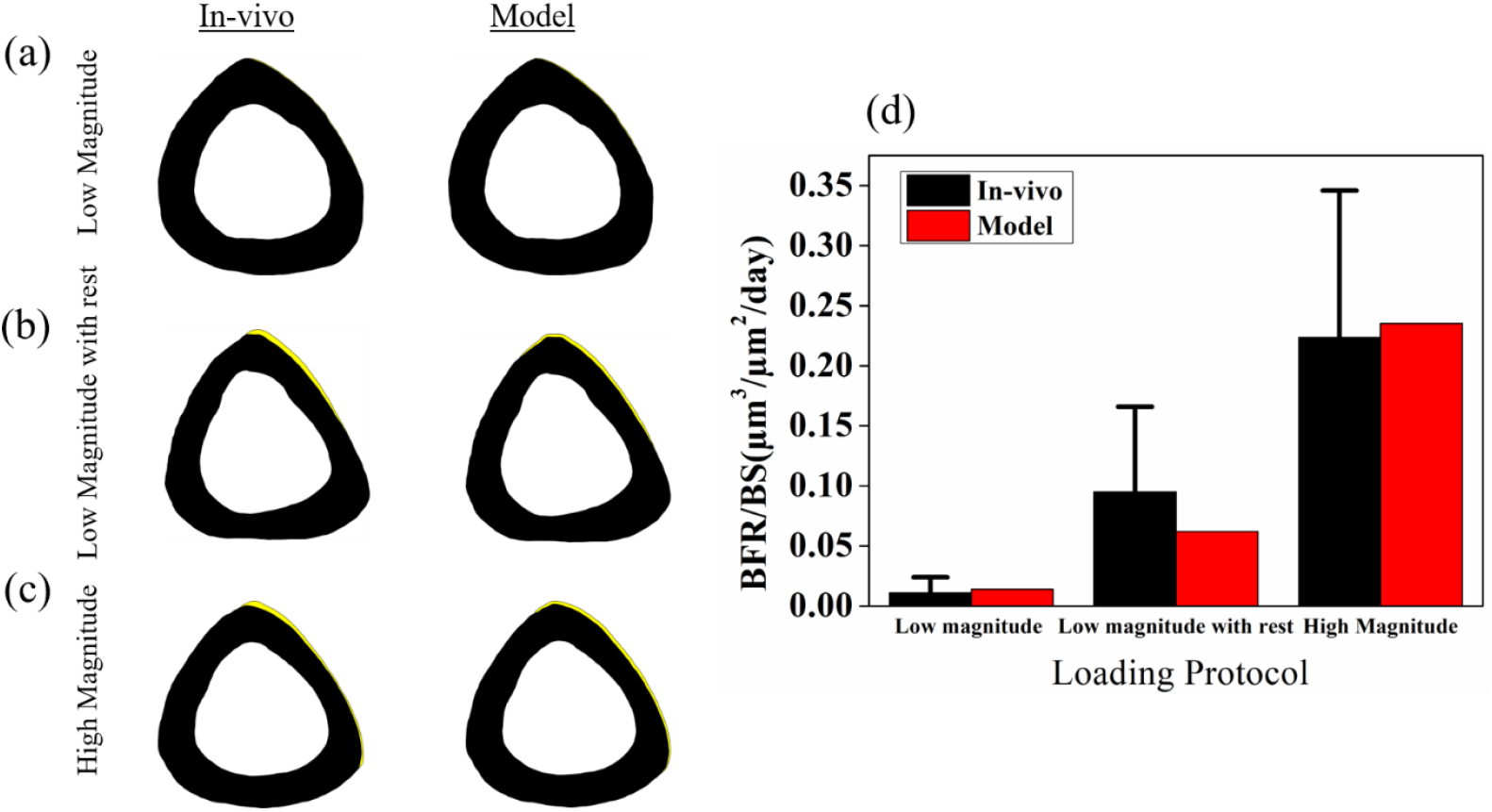
Comparison of experimental new bone formation with model predictions when dissipation energy density induced by the interaction of fluid velocity and pore pressure (*φ* = *φ*_*p*_ + *α*^2^*φ*_*v*_ − 2*αφ*_*pv*_) was utilized as the stimulus: (a) site-specific new bone formation under the low-magnitude loading protocol (yellow indicates new bone formation); (b) site-specific new bone formation under the low-magnitude loading protocol with rest insertion; (c) site-specific new bone formation under the high-magnitude loading protocol; (d) comparison of experimental and model-predicted BFR/BS values.

**Low Magnitude:** The same dissipation energy density (*φ* = *φ*_*p*_ + *α*^2^*φ*_*v*_ − 2*αφ*_*pv*_) was used as a stimulus to predict osteogenesis under the low-magnitude loading. A comparison of experimental and model-predicted BFR/BS values is shown in Fig. 10(d). Statistical analysis indicates no significant difference between the experimental BFR/BS (0.011±0.013 *μm*^3^/*μm*^2^/*day*) and the model-predicted BFR/BS (0.014 *μm*^3^/*μm*^2^/*day*, p=0.82, Student’s t-test). The site-specific bone distribution of new bone formation (Fig. 10(a)) was also compared using the Watson U^2^ test and was found not to differ significantly (p = 0.78, Watson U^2^ test).

**Low Magnitude with Rest Insertion:** Introducing a 10-second rest between loading cycles significantly enhanced new bone formation compared to the continuous low-magnitude loading protocol (Fig. 10(d)). The periosteal new bone formation induced by the rest-inserted protocol was evaluated using both the Student’s t-test and the Watson U^2^ test. The bar chart comparison showed that the model-predicted BFR/BS value (0.0621 *μm*^3^/*μm*^2^/*day*) was not significantly different from the experimental BFR/BS value (0.095 ± 0.071 *μm*^3^/*μm*^2^/*day* p = 0.86, Student’s t-test). The site-specific new bone formation (Fig. 10(b)) was also not significantly different (p=0.89, Watson U^2^ test).

### 3.4 Hypothesis Testing

The *p*-values for all the cases described above are summarized in Table 2.

**Table 2:** Comparison of *p*-values for the Student’s t-test and the Watson’s U^2^ test for all cases.

|  | High Magnitude |  | Low Magnitude |  | Low Magnitude with Rest Insertion |  |
| --- | --- | --- | --- | --- | --- | --- |
| | Student's t-test | Watson's $U^2$ test | Student's t-test | Watson's $U^2$ test | Student's t-test | Watson's $U^2$ test |
| $\varphi_p$ | 0.92 | 0.025 | 0.04 | 0.003 | 0.62 | $3.5 \times 10^{-5}$ |
| $\alpha^2 \varphi_v$ | 0.72 | 0.036 | 0.30 | 0.046 | 0.69 | 0.011 |
| $\varphi_p + \alpha^2 \varphi_v$ | 0.71 | 0.035 | 0.04 | 0.33 | 0.70 | 0.007 |
| $\varphi_p + \alpha^2 \varphi_v - 2\alpha\varphi_{pv}$ | 0.87 | 0.70 | 0.82 | 0.78 | 0.86 | 0.89 |

It is evident from Table 2 that *p* < 0.05 for Watson’s U^2^ test across all loading protocols (high magnitude, low magnitude, and low magnitude with rest insertion) for all stimulus cases except the interactive combination of pore pressure and fluid velocity (*φ*_*p*_ + *α*^2^*φ*_*v*_ − 2*αφ*_*pv*_). For this interactive case, *p* > 0.05 for both Student’s t-test and Watson’s U^2^ test. This outcome rejects the null hypothesis that none of the four dissipation energy cases can model the asymmetric osteogenesis. It supports the alternative hypothesis that the dissipation energy density based on the interaction of pore pressure and fluid velocity can accurately represent the experimental results.

## 4. Discussion

The study presents a robust methodology highlighting the role of load-induced pore pressure developed within the lacunar-canalicular network (LCN) in bone mechano-transduction. A mathematical model was developed to explain why cantilever-induced new bone formation occurs predominantly at the anterolateral side, despite the posteromedial side of the mid-diaphysis experiencing the same magnitude of peak strain, fluid velocity, and pore pressure. By integrating both load-induced fluid flow in the LCS and load-induced pore pressure, the model provides deeper insight into the mechano-transduction process of bone.

The developed model successfully predicted cantilever loading-induced osteogenesis under low-magnitude, low-magnitude with rest insertion, and high-magnitude protocols with spatial accuracy. It captured not only the site-specific distribution of new bone formation but also the bone formation rate per unit bone surface (BFR/BS) across all loading protocols.

Importantly, the mathematical framework is generalizable and can be applied to predict osteogenesis under any loading protocol, regardless of waveform. Because the present methodology employs Fourier series, dissipation energy density can be computed for arbitrary loading waveforms. This makes the model uniquely suited for comparing osteogenesis across different loading waveforms and for guiding the design of exercise regimens for bone regeneration.

Earlier literature proposed strain-based mathematical models to predict osteogenesis, correlating peak strain with periosteal new bone formation only [30] [32]. However, predicting osteogenesis on both periosteal and endosteal surfaces remained a challenge. In vitro studies revealed that osteocytes respond only at very high strain levels, about (∼30000 µε), while physiological loading typically induces tissue strains of ∼2000µε [33], which is not sufficient to trigger osteocytic release biochemical signals for osteogenesis. This limitation shifted attention towards load-induced fluid flow within the LCS. Munro and Knoth provided experimental evidence for fluid flow, while Weinbaum and colleagues established the mathematical foundation. [29] [21].

Further research into the osteocyte microenvironment introduced the concept of strain amplification [34], describing how tissue-level strain amplifies into the cellular-level strain [34]. When bone is mechanically loaded, a pressure gradient develops within the porosities of the LCN, driving fluid flow within the LCS. Osteocyte processes are anchored to the bone matrix via tethering fibers (Fig. 5 (c)). As fluid passes through pericellular spaces, the cell process deforms radially, leading to dissipation of energy, which has been assumed to stimulate osteocyte activation. This paradigm has motivated mathematical linking loading-induced stimuli to new bone formation.

Biot’s theory of poroelasticity has been widely applied to model cortical bone, typically assuming periosteal and endocortical surfaces to be impermeable and permeable, respectively. This permeability of the endocortical surface results in higher fluid velocity, while the periosteal surface, being impermeable, shows negligible fluid velocity. Thus, fluid flow alone cannot account for new bone formation observed simultaneously at both surfaces, promoting further investigation into the LCS.

Over the past five decades, the role of load-induced fluid flow in bone mechanotransduction has been well recognized [5] [6] [35] [36] [37] [38], whereas the role of load-induced pore pressure has been largely overlooked. Singh et al. revisited the mechanical environment of the LCS and showed that pore pressure, in addition to fluid flow, plays a critical role in mechanotransduction. Their mathematical model correctly predicted bone formation at both surfaces simultaneously [18]. Interestingly, new bone formation was found to localize at sites of elevated stimulus. However, Srinivasan et al.[20] reported a puzzling observation: in cantilever-loaded mouse tibia, new bone formation occurred predominantly at the anterolateral side of the mid-diaphysis, even though the opposing posteromedial surface experienced equal peak strain, fluid velocity, and pore pressure. This highlighted the need to examine the osteocyte microenvironment more closely.

While Singh et al.[18] considered pore pressure and fluid velocity without interaction-attributing fluid flow to endocortical bone formation and pore pressure to periosteal bone formation-the present study demonstrates that a complete picture emerges only when their combined effect with interplay is taken into account. Specifically, dissipation energy density arises from the interaction of these two stimuli at the osteocyte level. In this work, osteocytes and their processes were modeled as a viscoelastic material, and a methodology was proposed to compute the dissipation energy density, combining load-induced fluid velocity and pore pressure developed within LCS. The developed mathematical model successfully predicted in vivo data reported in the literature (Fig. 10), including site-specific distributions under multiple loading protocols.

A particularly important phenomenon captured by the model is the effect of rest insertion. Experimental data have shown that a 10-second rest between loading cycles significantly enhances new bone formation (Fig. 10(b)). The underlying mechanism can be explained by analysing fluid velocity and pore pressure dynamics (Fig. 11). At the onset of loading, pores are relatively empty, and the pressure gradient drives fluid flow. Under continuous loading, however, fluid becomes trapped, and velocity reaches a steady state (Fig. 11(a)). In contrast, when rest intervals are introduced, fluid has sufficient time to escape and re-enter the pores, resulting in higher fluid velocities and greater pore pressure fluctuations in subsequent cycles (Fig. 11(b)). This dynamics enhances dissipation energy and, consequently, osteogenesis.

**Figure. 11:**
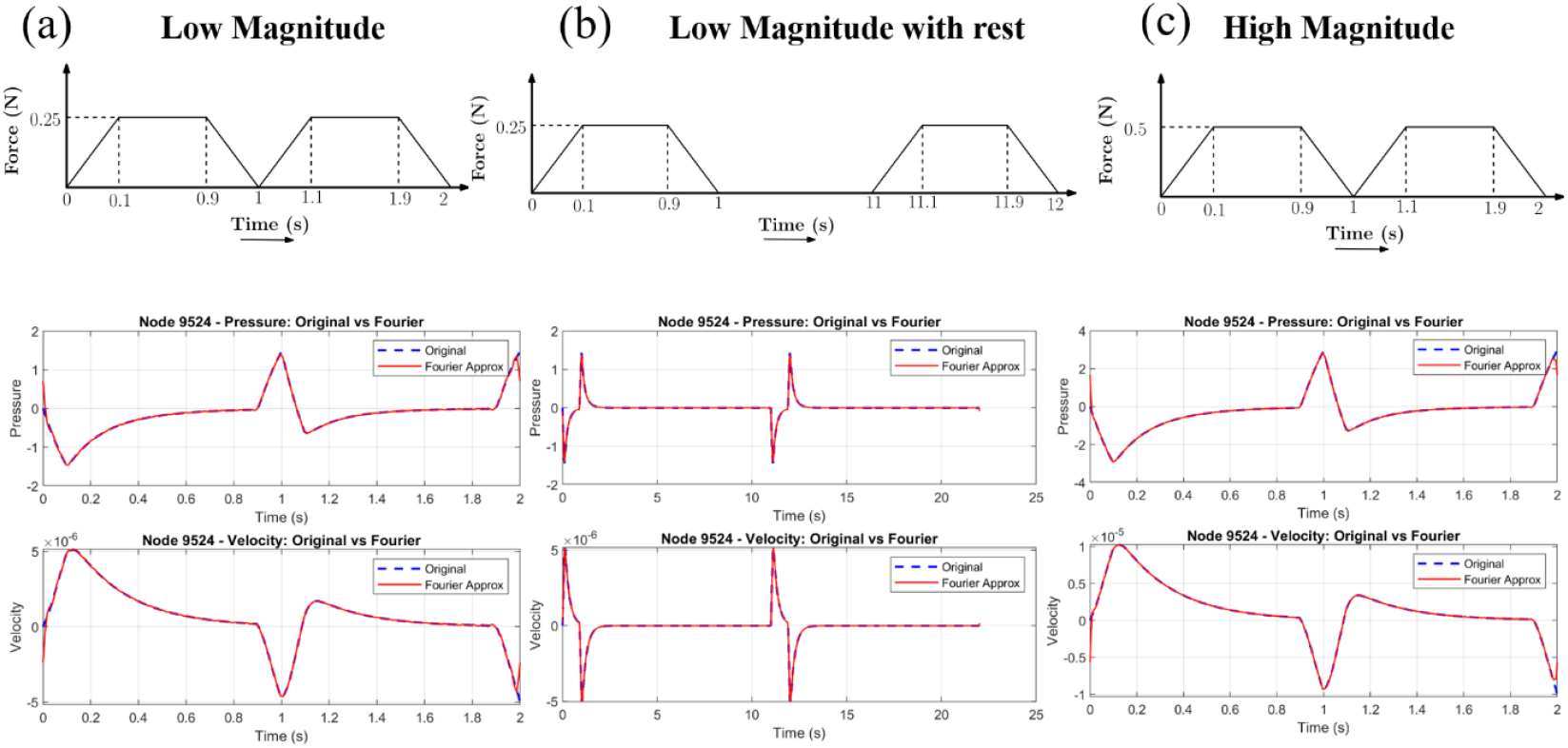
Variation of fluid velocity and pore pressure with respect to time at node 9524 (tensile side of the bone cross-section). The time-dependent pore-pressure and fluid velocity were approximated using a Fourier series for the respective loading protocols: (a) low magnitude, (b) low magnitude with rest insertion, and (c) high magnitude.

### Summary of the present work

- Load-induced fluid flow and pore pressure within the lacunar–canalicular system (LCS) are essential mechanotransductive stimuli; however, it is their combined interaction that governs new bone formation.
- The interaction of these stimuli successfully predicted the site of new bone formation and explained why the tensile side of the bone cross-section is more susceptible to osteogenesis.
- The model accurately reproduced experimental observations across three groups of mice and three different loading protocols, highlighting its robustness.
- A key contribution of this work is the development of a Fourier series–based methodology to compute dissipation energy density. This generalizable approach enables the computation of stimuli under any arbitrary loading waveform and offers potential for systematically comparing osteogenic responses across diverse loading regimes.

The present model was developed with certain simplifications, which are listed below:

- The model does not incorporate the complete cellular environment, such as the attachment of tethering fibres to the bone matrix or the role of integrins.
- Bone was idealized as a linear, isotropic poroelastic material, whereas cortical bone in reality exhibits anisotropy.
- Only lacunar–canalicular (LCN) porosity was considered for fluid flow analysis. Since cortical bone contains porosities at multiple length scales (e.g., LCN and vascular porosities), incorporating vascular porosity alongside LCN may further strengthen the model.

## 5. Conclusions

This study establishes that the interaction between load-induced pore pressure and fluid flow within the lacunar-canalicular system (LCS) governs osteogenesis more accurately than either stimulus alone. A poroelastic finite element model was developed to quantify this interaction and predict the spatial distribution of new bone formation under cantilever loading. The model was validated against in vivo data for multiple loading protocols – low magnitude, low magnitude with rest insertion, and high magnitude – and demonstrated close agreement in both site-specific new bone distribution and bone formation rate per unit bone surface (BFR/BS). Statistical analyses using Student’s t-test and Watson’s U^*2*^ test confirmed that only the interactive stimulus accurately reproduced experimental observations.

Unlike previous models that depend on the specific shape of the loading waveform, the present approach uses a Fourier series-based formulation to compute dissipation energy density, enabling analysis under arbitrary loading conditions. This methodological generality allows for systematic comparison of osteogenic responses across diverse mechanical stimuli and facilitates the design of optimized loading or exercise regimens for enhancing bone formation and preventing bone loss.

To conclude, the model not only elucidates why osteogenesis localizes predominantly on the tensile side despite equivalent strain magnitudes at the compressive side but also provides a unifying mechanistic framework for future investigations into bone mechanotransduction. Its capacity to integrate complex mechanical interactions into a single quantifiable parameter represents a significant step toward predictive modeling of bone adaptation and clinically relevant therapeutic strategies.

## Contributions

**H.S**. was involved in conceptualization, mathematical modeling, data collection, optimization, model validation, statistical analysis, software development, manuscript writing, and editing. **J.P**. supervised and conceptualized the study, contributed to the model review, and edited the manuscript.

## Data availability

All data supporting the findings of this study are available within the article.

## Ethics declarations

### Conflict of interest

The authors declare that they have no conflict of interest.

## Notes

### Competing Interest Statement

The authors have declared no competing interest.

### Summary of Updates

Additional data and figures have been incorporated to strengthen the credibility of the results. Accordingly, the abstract, title, results, and conclusions have been revised to make the study more comprehensive.

